# High throughput CRISPR screening identifies genes involved in macrophage viability and inflammatory pathways

**DOI:** 10.1101/807164

**Authors:** Sergio Covarrubias, Apple Vollmers, Allyson Capili, Michael Boettcher, Elektra K. Robinson, Laura O’Briain, Christopher Vollmers, James Blau, Michael McManus, Susan Carpenter

## Abstract

Macrophages are critical cells of the innate immune system involved in the recognition and destruction of invading microbes in addition to the resolution of inflammation and maintenance of homeostasis. Understanding the genes involved in all aspects of macrophage biology is essential to gaining new insights into immune system dysregulation during diseases that range from autoinflammatory to cancer. Here we utilize high throughput clustered regularly interspaced short palindromic repeats (CRISPR) screening to generate a resource that identifies genes required for macrophage viability and function. First, we employ a pooled based CRISPR/Cas nuclease active screening approach to identify essential genes required for macrophage viability by targeting genes within coding exons. In addition, we also target 3’UTRs to gain insights into new *cis*-regulatory regions that control expression of these essential genes. Second, using our recently generated NF-κB reporter macrophage line, we perform a fluorescence-activated cell sorting (FACS)-based high-throughput genetic screen to identify regulators of inflammation. We identify a number of novel positive and negative regulators of the NF-κB pathway as well as unraveling complexities of the TNF signaling cascade showing it can function in an autocrine manner to negatively regulate the pathway. Utilizing a single complex library design we are capable of interrogating various aspects of macrophage biology, generating a resource for future studies.

**Significance:** Excess inflammation is associated with a variety of autoimmune diseases and cancers. Macrophages are important mediators of this inflammatory response. Defining the genes involved in their viability and effector function is needed to completely understand these two important aspects of macrophage biology. Here we screened over 21,000 genes and generated a resource guide of genes required for macrophage viability as well as novel positive and negative regulators of NF-κB signaling. We reveal important regulatory aspects of TNF signaling and showing that membrane-bound TNF primarily functions in an autocrine fashion to negatively regulate inflammation.

## Introduction

Clustered regularly interspaced short palindromic repeats (CRISPR) technology has revolutionized the field of functional genomics providing an easy-to-use method for disrupting specific genes (1). The coupling of CRISPR technology with pooled sgRNA screening allows simultaneous knockout of thousands of individual genes in a large population of cells (2,3). Numerous CRISPR screens have probed pathways ranging from cell viability (4,5) to virus infection (6,7), supporting the use of this system for exploring a wide-range of biology. Our study utilizes CRISPR screening to answer three questions about macrophage biology: 1) What genes are required for macrophage survival and proliferation? 2) How are those genes regulated? 3) What genes contribute to the downstream inflammatory signaling processes? These screens provide a wealth of insight into macrophage function and will serve as a resource for future work aimed at better understanding the molecular mechanisms of action of the identified hits.

Macrophages are critical cells of the innate immune system providing one of the first lines of defense against invading microbes. Macrophages arise from precursor monocyte cells that constitute ~10-20% of the immune cells found in the blood (8). Upon encountering a danger signal, monocytes differentiate into macrophages and rapidly move to the site of infection. Important aspects of macrophage function include their ability to proliferate and migrate, as well as their ability to induce the inflammatory program to aid in clearing infections and initiate tissue repair to maintain homeostasis (9).

Here we performed two CRISPR screens to investigate macrophage biology. The first screen aimed to identify and characterize the regulation of genes essential to macrophage viability. Identification of the full catalog of essential genes is critical for obtaining a complete picture of the pathways important for a specific cell type (4). Cell-type specific differences in expression and splicing of genes required for viability may contribute to differential dependency on these genes across different cell lineages (4). Indeed, numerous viability screens have been performed in a wide-range of cell types and have demonstrated that different cell types rely on distinct pathways for survival (3,10,11). To this end, we built a complex CRISPR library targeting all protein coding genes and microRNAs in order to define all genes that are essential specifically for macrophage viability. In addition to identifying essential genes in macrophages we also aimed to understand what regulatory mechanisms could be involved in controlling expression of these genes. We know that the untranslated regions (UTRs) of messages are important for regulating stability, translation and localization (reviewed in (12)). Sequences within the UTRs (*cis*-elements) can function in microRNA binding, structure and/or protein binding (13). Furthermore, cell-type specific expression differences in microRNAs and RNA binding proteins (RBP) allows for specificity in regulating mRNAs (14). Given the rich source of regulation that occurs within the 3’UTR, we built into our library design sgRNAs targeting 3’UTRs within known essential genes with the goal of systematically identifying *cis*-elements that contribute to the regulation of essential genes.

The second screen aimed to identify genes involved in mediating inflammation, which is one of the key roles of macrophages. The inflammatory response is initiated through recognition of specific pathogen-associated molecular patterns (PAMPs) through their germline-encoded pattern recognition receptors (PRRs) (15). These receptors couple pathogen-sensing to activation of downstream signaling cascades resulting in activation of numerous transcription factors, including NF-kappa B (NF-κB) (16,17). NF-κB is a major transcription factor that drives the inflammatory response (18,19). We previously developed an immortalized bone marrow-derived macrophage line (iBMDM) with CRISPR editing capabilities and an NF-κB-responsive GFP-fluorescence reporter that allows monitoring of the NF-κB activation state of the cell (20). Our iBMDM-NF-κB-Cas cells are the first immune-relevant NF-κB -reporter cells that are fully compatible with FACS-based CRISPR-pooled screening enabling us to use our pooled screening approach to identify both positive and negative regulators of inflammation.

Together, using these two screens, we probed three different parameters related to macrophage biology. We provide a resource that identifies genes essential to macrophage viability and provide insights into potential *cis*-elements within 3’UTRs of these genes that may reveal important means of regulation of said genes. Lastly, we identify novel positive and negative regulators of NF-κB inflammatory signaling. Unexpectedly from our screen we uncover a role for tumor necrosis factor (Tnf) as a strong negative regulator of NF-κB and show this is functioning in an autocrine manner. In a single screen, we bring together decades of literature on the complex regulation of Tnf. Here we demonstrate the power of CRISPR pooled screening to identify a plethora of genes with varied and critical roles in macrophage biology.

## Results

### Pooled CRISPR Screen identifies macrophage-specific genes involved in viability

In order to define all genes essential for macrophage survival, iBMDM-Cas9 cells were transduced (MOI=0.3) with pooled lentivirus generated from our custom whole-genome sgRNA library containing ~270K individual sgRNAs targeting all RefSeq annotated coding genes and microRNAs (12 guides/gene), along with ~5K non-targeting controls (Figure 1A, S1A, Table S1). We obtained an initial infection coverage of >200 cells/sgRNA and maintained cells at >1000 cells/sgRNA throughout the screen. Cells were cultured for 20 days collecting genomic DNA from cells at Day 0 and Day 20 (Figure 1A). The libraries were prepared as described previously (21). Using the Mann-Whitney (MW) U test, we compared the sgRNA repertoire of Day 20 to Day 0 and identified significant genes (Figure 1B). We identified expected viability-related genes with roles in spliceosome, proteasome and cell cycle functions (Figure 1C). The majority of the top significant hits were genes essential for viability, while only 1% of genes were growth suppressors, consistent with previous findings (Figure 1D, (22)). We compared our viability screen hits to the GenomeCRISPR (www.genomecrispr.org/) database, which includes a collection of ~500 CRISPR viability screens performed in ~421 cell types (Figure 1E). Over 93% of genes from our screen overlapped those from the database (Figure 1E) suggesting that these are genes critical for a variety of biological processes common to all cell types. Interestingly, ~6% of genes identified showed the opposite MW Z-score phenotype in our screen compared to the database suggesting that these genes have unique cell-type-specific functions in macrophages (Figure 1E). Furthermore, these macrophage-specific essential hits included genes involved in NF-κB signaling (Figure S1) suggesting that these genes evolved functions involving not only survival but also in inflammatory activation, a critical component of macrophage function.

**Figure 1:**
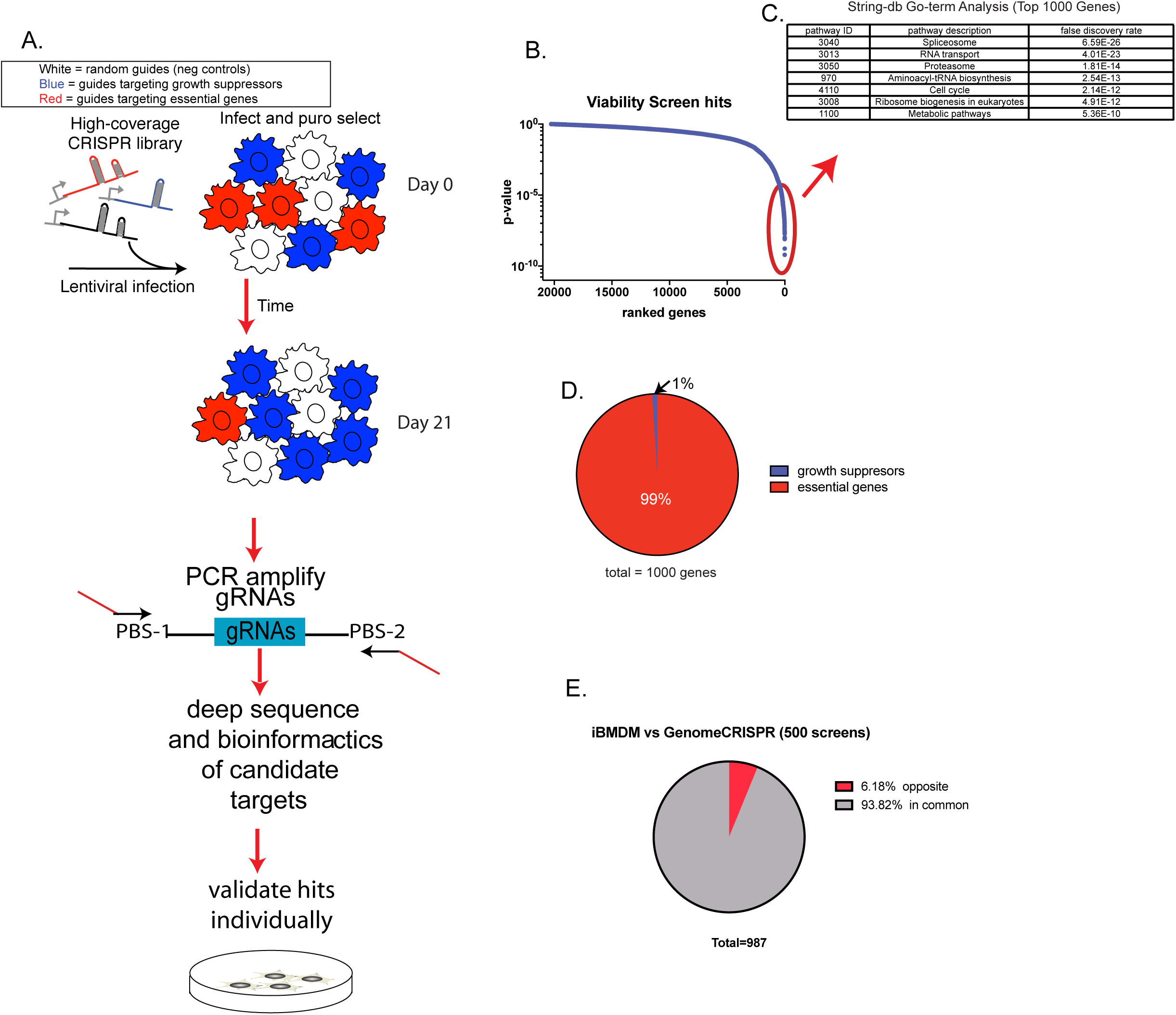
Screen identifies macrophage-specific genes involved in viability. A. Cas9-expressing iBMDM cells were infected with a whole-genome library targeting all refseq annotated coding genes (S1 Table). Two days post infected, cells were harvested to collect the day 0 time-point and another time-point was collected after day 20 in culture. B. The Mann-Whitney U test was performed comparing 12 sgRNA targeting each gene to the nontargeting controls for both Day 20 and Day 0 collected samples and significant genes are displayed ranked by significance. C. Go-term analysis was assessed for the top 1000 significant hits using String-DB. D. We determined the number of essential genes (negative MW Z-score) and total number of growth suppressor genes (positive MW Z-score) and plotted them as a fraction of total (total=1000 genes). E. Viability screen hits from our screen were compared to those from GenomeCRISPR, a collection of ~500 CRISPR screens (www.genomecrispr.org/). Genes “in common” as well as genes showing “opposite” phenotypes are displayed.

### CRISPR Targeting of the 3’ UTRs of essential genes identifies novel *cis* regulatory elements

UTRs offer a critical source of regulation for messages through specific *cis*-elements, which can bind microRNAs and/or proteins to regulate pathways including RNA decay and translation (12). To probe for novel *cis*-elements within 3’UTRs, we specifically targeted 3’UTR within known essential genes. For these essential genes, we expected that guides targeting coding exons would cause a decrease in fitness. In contrast guides targeting 3’UTR cis-elements that result in an increase in fitness will represent regions containing regulatory *cis*-elements with negative roles on gene expression (microRNA binding sites, etc.). We assessed the phenotypes of 3’UTR-targeting sgRNAs and found an overall neutral average phenotype for these guides (Figure 2A) suggesting that the majority of the sites we targeted did not contain any *cis* regulatory elements. However, a subset of these 3’UTR-targeting guides demonstrated phenotypes of >3-fold change in both directions suggestive of sites that contain regulatory elements that could improve or decrease fitness of the cells (Figure 2B-D). We focused on 3’UTR guides that showed positive (>3-fold) enrichment for genes whose coding-targeting guides demonstrated significant negative enrichment (Figure 2B-D). We reasoned that these 3’UTR-targeting guides may disrupt significant *cis*-elements that help explain the opposing phenotype we observed (Figure 2B-D). We individually cloned selected candidate 3’UTR guides and validated the phenotype using a mixcell proliferation assay (Figure 2E). The mix cell assay involved combining cherry positive cells (containing guide RNAs) with cherry negative cells at a 1:1 ratio and monitoring cell growth over time as assessed by changes in the ratio of cherry positive to cherry negative cells. In all the selected hits we could validate that indeed the elements we targeted resulted in an increase in fitness compared to targeting the coding sequence which resulted in decreased fitness. The phenotypes for these 3’UTR-targeting guides provide a resource of potentially important *cis*-elements involved in mRNA stability and further work could involve interrogating whether these sites are microRNA targets or targets of other proteins involved in RNA decay or translation processes.

**Figure 2:**
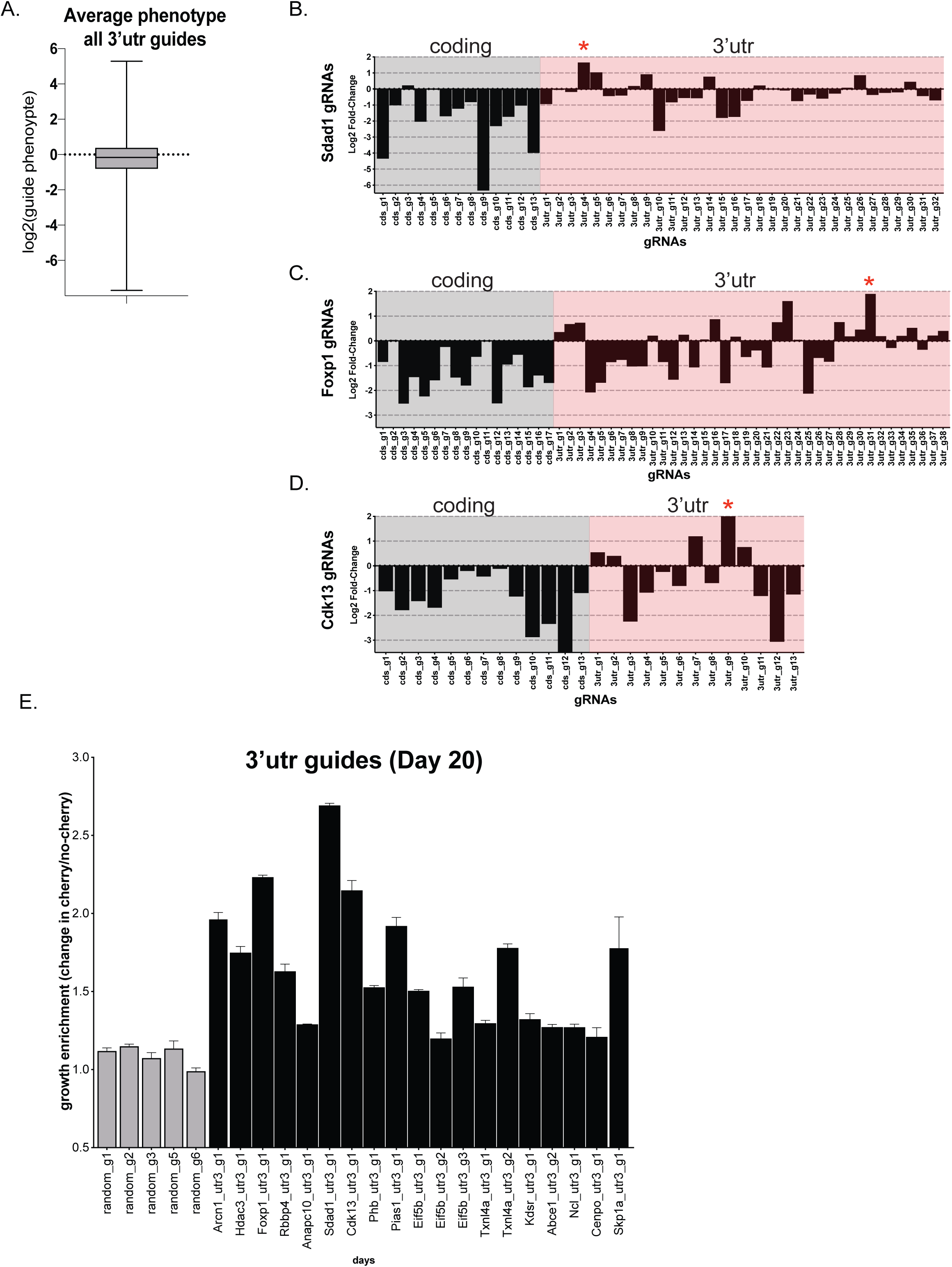
Targeting of the 3’ UTRs of essential genes to probe for novel cis regulatory elements. A. Average phenotypes for all 3’utr-targeting guides is plotted. B-D. We summarize the phenotypes for coding-targeting guides (gray) and 3’utr-targeting guides (pink) for select genes: Sdad1, Foxp1 and Cdk13. E. We selected 3’utr-targeting guides that demonstrated positive enrichment, opposite to their coding-targeting guides, which had negative enrichment. We validated select guides by monitoring growth of sgRNA infected cells (cherry) relative to an uninfected reference cells in a mix-cell growth assay.

### FACS-based reporter screen identifies novel regulators of NF-κB

Macrophages are critical effectors of the inflammatory response, which involves transcription factors including NF-κB (18). We had previously developed an NF-κB responsive GFP-reporter system in iBMDMs (iBMDM-NF-κB-Cas) and demonstrated lipopolysaccharide (LPS)-dependent activation of GFP-fluorescence (20). Here we performed a FACS-based sorting screen using iBMDM-NF-κB-Cas cells and infected with the same library as outlined in Figure 1. After the library was established in the cells for 7 days, we stimulated with LPS for 24h (Figure 3A, S2A-B). We sorted the top/bottom 20% of GFP expressing cells and collected approximately 100 cells per sgRNA (~27 million cells for each top/bottom sort) with the aim of identifying both positive (bottom 20%) and negative regulators (top 20%) of the pathway (Figure 3F, S2A-B). We performed a Mann-Whitney U test to identify significant genes, comparing GFP-low to GFP-high sorted samples, and significant genes were ranked by p value (Figure 3B-C). As expected, we found several positive controls including eGFP and known regulators: Myd88 and Rela in our top hits (Figure 3C). We performed go-term enrichment analysis for the top 150 significant positive regulators and found enrichment for pathways that included “NF-kappa B signaling” (Figure 3D). Within the top 150 genes, we identified numerous genes known to be involved in the TLR/NF-κB signaling pathway, which include Tlr4 (LPS receptor) and Rela (NF-κB) (Figure 3E), confirming that the screen was a success. We plotted the average phenotypes (top 3 guides) for our top 40 candidates, which showed that the average sgRNA-enrichments were significant and could be validated (Figure 3G). Top candidates were localized throughout the cell (Figure 3H), including in the extracellular compartment. Numerous positive and negative regulators of NF-κB have been identified by their differential expression upon NF-κB activation (23). Using previously published data (24), we examined the top 50 negative and positive regulators and found that the majority were not differentially expressed during LPS stimulation and therefore could have been missed by previous approaches as regulators of the pathway (Figure S2C). NF-κB has been demonstrated to activate genes that function in positive or negative feedback regulation of the pathway (25). We assessed whether NF-κB (p65) bound to the promoters of our top candidates using published p65 Chip-seq data (26)(Figure S2D-E). We found that p65 bound 42% and 54% of the top positive and negative regulators respectably (Figure S2D-E) further supporting the idea that a significant number of regulators of NF-κB can be themselves regulated by NF-κB. We validated our top candidates by re-cloning the top two performing sgRNAs per candidate, generating individual cell lines for each sgRNA, followed by lentiviral infection, selection and LPS stimulation for 6h (Figure 3I). The readout for our secondary validation experiments involved measuring Il6 by qPCR. Il6 is a well-known downstream target of NF-κB and a crucial gene in controlling inflammation (Figure 3I). As expected, ablation of our negative regulators resulted in increased Il6 expression, while targeting of positive regulators led to decreased levels of Il6 relative to non-targeting controls (Figure 3I). In summary, we performed a genome-wide screen and identified 50 novel positive and 65 negative regulators of NF-κB signaling. To date there are 120 known regulators of NF-κB and our screen and subsequent validation has added 115 new regulators (p value <0.01) to the pathway. These will provide a rich source of information going forward to better understand the complex pathways that control inflammation.

**Figure 3:**
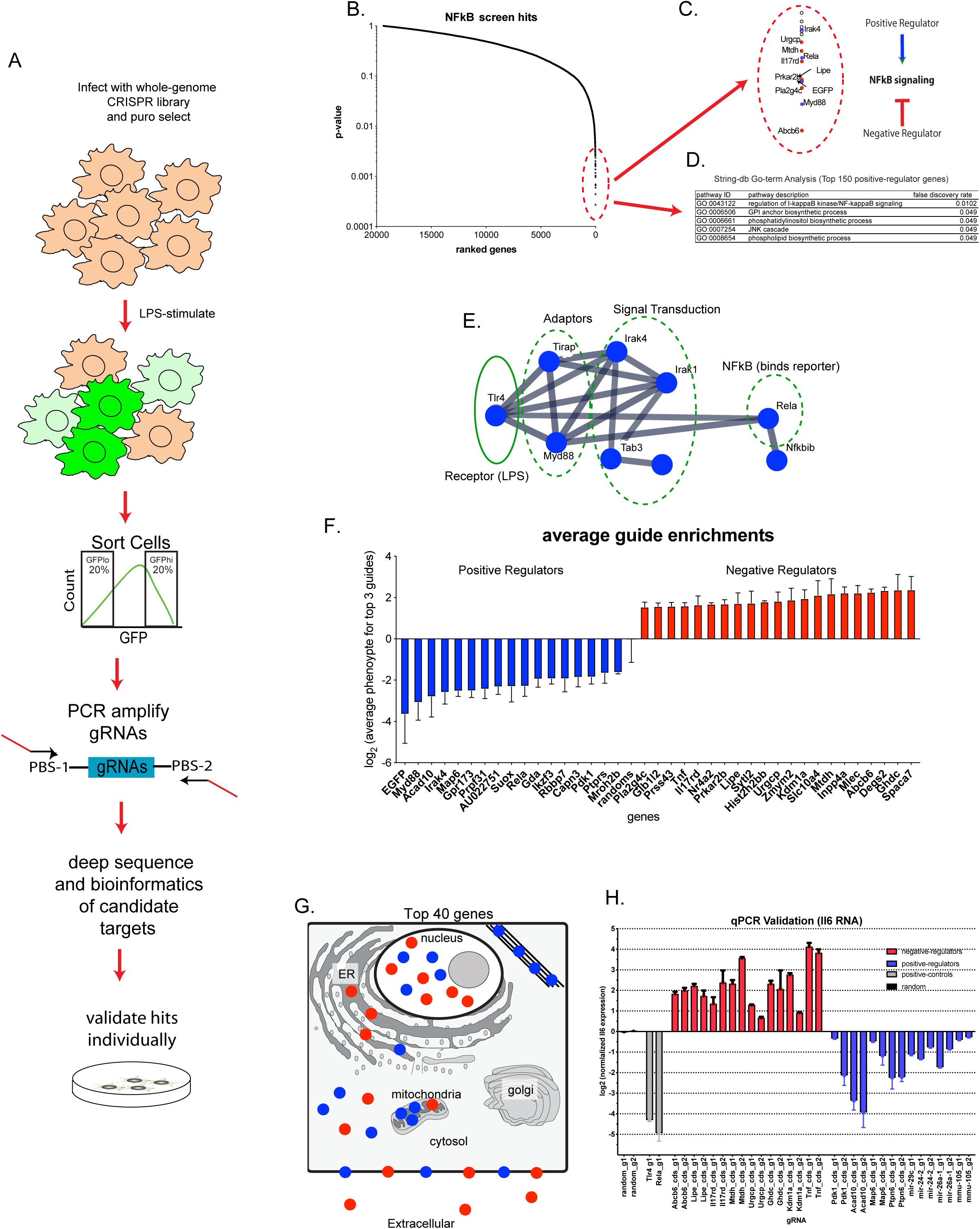
Screen Identifies Novel Positive and Negative Regulators of NFKB signaling. A. Overview of the NFKB screen: sgRNA library-infected iBMDM-NFKB-Cas9 cells were stimulated with LPS (200ng/ml) for 24h prior to sorting the top and bottom 20% of GFP-expressing cells. Cells were collected and processed as described in methods. B. Mann-Whitney U test was performed comparing 12 sgRNA targeting each gene to the nontargeting controls for GFP-low vs GFP-high sorted samples and significant genes are displayed ranked by significance. C. Zoom in of the top screen hits, displaying positive regulator in blue and negative regulators in red and Diagram depicting positive/negative regulation NFKB signaling. D. Go-term analysis was assessed for the top 150 positive regulators using String-DB. E. Connectively was determine by stringDB for the top 150 positive regulator candidates. F. Average sgRNA enrichment for the top 3 sgRNA was calculated for the top 40 screen hits. G. Predicted protein localization was determined for the top 40 most significant genes using uniprot’s compartments DB (https://compartments.jensenlab.org/Downloads). H. Selected candidates were infected with either control (random) or candidate-specific sgRNAs and were stimulated for 6h with LPS, prior to RNA-harvest. qPCR-based validation was performed by performing qRT-PCR for Il6 RNA relative to Gapdh. Experiment was repeated 3 times and a representative experiment is displayed.

### Membrane bound tumor necrosis factor alpha (Tnf) acts as a strong negative regulator of NF-κB pathway

The tumor necrosis factor alpha (Tnf) is a well-known pro-inflammatory cytokine with established roles in driving NF-κB-related inflammation (27). Furthermore, anti-Tnf therapy is a proven method for the treatment of various inflammatory-related diseases (28). Given its established role as a soluble protein functioning as a positive regulator of NF-κB, it was surprising to discover our pooled based approach screen identified Tnf as a strong negative regulator of inflammation (Figure 3I). We confirmed Tnf-editing by stimulating control or anti-Tnf edited cells with LPS for 24h prior to collecting supernatant and analyzing Tnf protein via ELISA analysis (Figure 4A). We found Tnf was undetectable in the supernatant following LPS stimulation (Figure 4A). We also confirmed near complete ablation of Tnf via intracellular-staining (Figure 4B). Our screen suggested a localized function for Tnf, which would be incompatible with its soluble state. Interestingly, Tnf can exist as both soluble and membrane-bound (29). We confirmed the presence of membrane-bound Tnf on the surface of our iBMDMs and found maximal surface Tnf at 6h post LPS stimulation (Figure 4C). Tnf mediates its inflammatory affect via binding to its receptor Tnfrsf1a (p55) (27). Our screen confirmed Tnfrsf1a (p55) as a positive regulator of NF-κB in contrast to what was found for Tnf (Figure 4D-E). However, Tnf can bind to two different receptors: Tnfrsf1a (p55) and Tnfrsf1b (p75), which may have opposing functions (30). Interestingly, following 6h post LPS stimulation, levels of Tnf and Tnfrsf1b (p75) increased 11-fold and 66-fold respectably, while the levels of Tnfrsf1a (p55) increase a moderate 4-fold (Figure 4F). We evaluated whether the enhanced activation of NF-κB in Tnf-edited cells could be rescued by mixing these cells with unedited (Tnf-expressing) cells. We combined cherry positive cells (containing Tnf guide RNAs) with unedited cherry negative cells at a 1:1 ratio. Mixed cells were LPS-stimulated for 24h prior to FACS analysis to measure GFP MFI (NF-κB activation) (Figure 4G). The enhanced NF-κB activation in Tnf-edited cells could not be rescued by mixing these cells with unedited cells (even when we increased the cherry negative cells to >75%). These data suggest that Tnf has autocrine properties, acting within the cell from which it is produced and neither soluble TNF production or membrane bound TNF from neighboring cells can reverse the increase in inflammatory signaling (Figure 4G-H). The expression profiles of TNF, TNFRSF1A and TNFRSF1B were mirrored in the human THP1 monocytic cell line stimulated with PAM3CSK4 (TLR1/2 agonist), suggesting this specific regulation of TNF is conserved (Figure S3A). Here we confirm that indeed it is membrane bound Tnf that is functioning as a negative regulator of NF-κB presumably through interactions with p75. Remarkably our single screening assay could provide insight into this complex signaling cascade and bring together decades of different approaches to reveal the complexity of Tnf signaling in macrophages involving membrane bound forms of Tnf in addition to the roles of the respective receptors.

**Figure 4:**
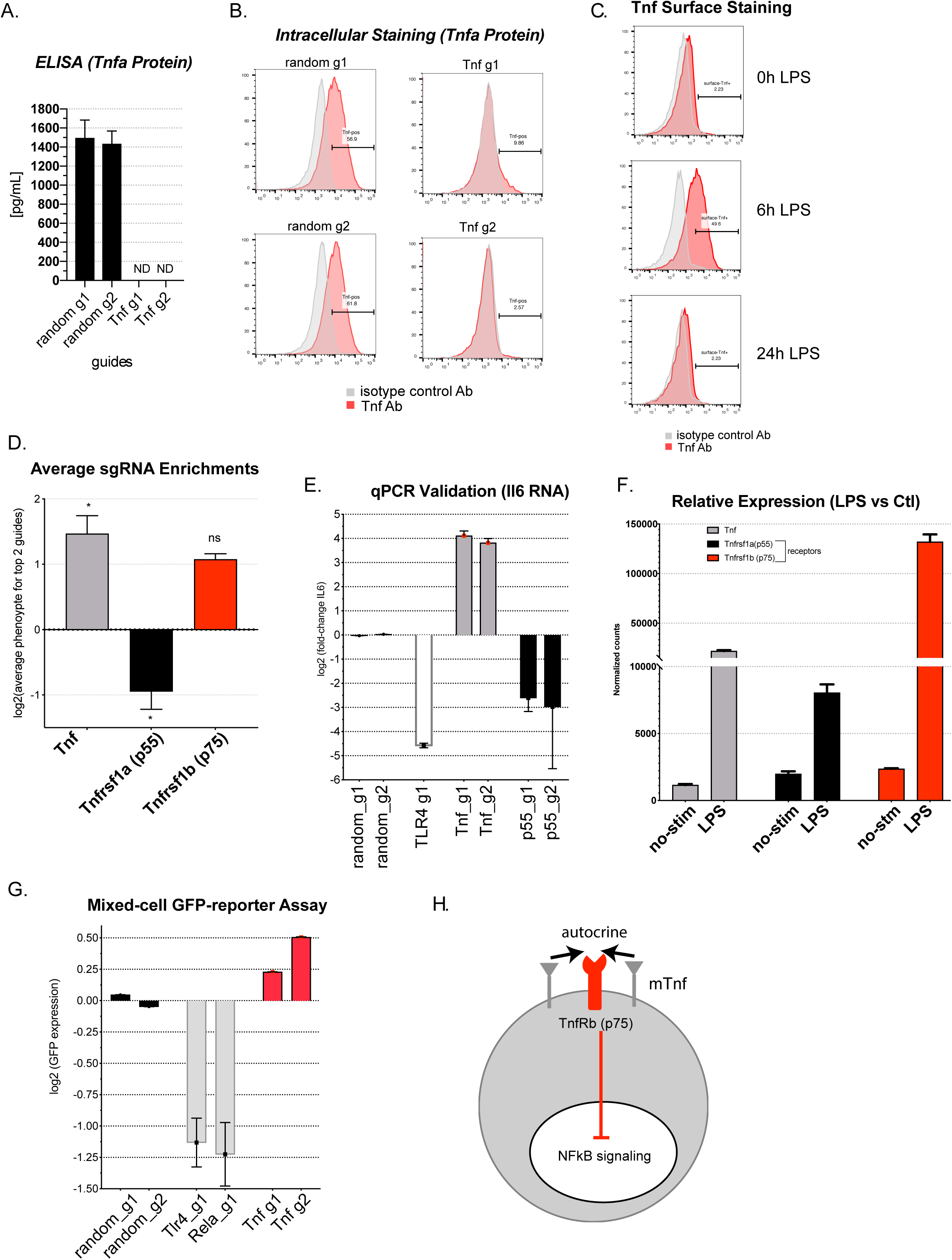
Screening identifies negative feedback regulatory loop used by the Tumor necrosis factor (Tnf) to regulate inflammation. A. ELISA was performed by first stimulating control or Tnf-edited iBMDMs for 24h with LPS, harvesting supernatants and quantitating levels of Tnf. B. Control or Tnf-edited iBMDMs were stimulated with LPS followed by Brefeldin A treatment and were then intracellular-stained to assess Tnf levels C. Average sgRNA enrichment for the top 2 sgRNAs was calculated for Tnf and its receptors: p55 and p75. D. qRT-PCR was performed for Il6 RNA in iBMDMs edited with either control, Tnf or p55 sgRNAs and stimulated for 6h with LPS. E. Mean FPKM values (4 biological replicates) are plotted for genes: Tnf, Tnfrsf1a and Tnfrsf1b for both un-stimulated (“no-stim”) or 24h LPS-stimulated cells. F. iBMDMs were stimulated with LPS for 0, 6, or 24h and were collected and surface-expression of Tnf was assessed G. Model of localized Tnf negative regulation of NFKB-signaling.

## Discussion

Here we utilize CRISPR-based pooled genetic screening to reveal genes important for both macrophage viability and inflammation. For the viability screen, we identified macrophage-specific viability genes, enriched in NF-κB -signaling (Figure S1A).

Additionally, we identified and validated viability phenotypes for 3’UTR-targeting guides, which may reveal important *cis*-elements involved in mRNA stability (discussed further below). In a separate screen, we utilized our NF-κB -reporter cells to perform a FACS-based screen resulting in the identification of novel positive and negative regulators of NF-κB signaling. We go on to describe an unexpected role for Tnf as a locally-acting negative regulator of NF-κB.

### Macrophage-specific genes involved in viability

Numerous viability screens have been performed in a wide-range of cell types and have demonstrated that the catalog of essential genes can vary significantly among distinct cell types (3,10). Cell-type specific differences in viability can be due to genetic differences that could render some pathways inactive, forcing dependency on other pathways. Indeed, Wang *el al* compared essential genes across 14 Acute Myeloid Leukemia (AML) lines and found certain genes to be essential only for specific subset of the AML lines with certain genotypes (10). Moreover, a typical cell expresses approximately two-thirds of its genes (31). Therefore, whether a gene or set of genes is essential can be context specific, varying upon different growth conditions or treatments etc. (32,33). In our screen, we identified 61 genes with distinct essentiality in macrophages compared to a database collection of ~421 cell lines CRISPR screens (Figure 1E, S1B). Not surprisingly, we identified the myeloid-specific transcription factor Interferon regulatory factor 8 (Irf8) as a macrophage-specific essential gene (Figure S1B). Interestingly, Irf8 has roles in both myeloid cell viability and inflammatory response (34). Additionally, the macrophage-specific essential genes included ones involved in NF-κB signaling suggesting that this pathway, similar to Irf8, may be important for both viability and inflammatory activation (Figure S1B). Indeed a significant fraction of them (~10 proteins), including Syk, Lyn are predicted to physically interact supporting a macrophage-specific regulation of protein complexes that control cell viability. In summary, understanding the pathways that control viability in macrophages is essential for development of novel gene targets to control macrophage proliferation in scenarios where the inflammatory response is not properly controlled.

### 3’UTR-targeting guides and identification of potential cis-elements

Untranslated regions (UTRs) of messages are important for regulating stability, translation and localization (12). Sequences within the UTRs (*cis*-elements) can function in microRNA binding, structure and/or protein binding (13). Here we targeted 3’UTRs of predominantly essential genes. For these genes, sgRNAs targeting coding sequences resulted in decreased fitness, as expected. Within these genes, we focused on 3’UTR-sgRNAs that resulted in increased fitness, which could represent disruption of destabilizing *cis*-elements, such as microRNA binding sites or AU-rich elements. CRISPR targeting of these destabilizing *cis*-elements may result in stabilization of the mRNA. We determined if there were any overlapping microRNA target sites using TargetScan, but did not find any with significant overlap suggesting that there are other mechanisms of regulation at play besides miRNA targeting. Although, Cdk13-utr3-g1 and Pias1-utr3-g1 (Figure 2E) did target regions near (<50-80bp) the microRNA binding sites for mir-124 and mir-10 respectably (Table S12). Another 3’UTR-targeting guide 1 for Arcn1, is predicted to target near a mushashi element, which is known to negatively regulate RNA stability (35). Given that the majority of our 3’UTR targeting guides resulted in no phenotype, an alternative method to try in future studies would be to use 2 sgRNAs to create larger deletions (36). In summary, we have functionally validated potential 3’UTR *cis*-elements and provide a rich resource for future work aimed at dissecting how these elements contribute to gene expression.

### Dissection the complex biology of the tumor necrosis factor (Tnf)

One of the surprising findings from our NF-κB screen was the identification of Tnf as a negative regulator of NF-κB (anti-inflammatory). We were surprised for two reasons: 1) The tumor necrosis factor (Tnf) is an extensively studied pro-inflammatory cytokine with established roles in driving NF-κB-related inflammation (27). 2) Tnf is a secreted protein and therefore a pooled CRISPR screen would not be expected to capture its biology. Numerous studies spanning decades of research have revealed that Tnf biology is much more complex. Tnf can exist as both soluble (17kD) and membrane-bound (26kD) forms and multiple groups have shown that membrane-bound Tnf functions as a negative regulator of inflammation in contrast to its soluble form (29,37). The dual functions of Tnf can also be explained in part by its binding to two receptors: Tnfrsf1a (p55) and Tnfrsf1b (p75), which have opposing effects on inflammation (30). Interestingly, the membrane-bound form of Tnf has been shown to preferably bind to the inhibitory receptor: p75, allowing for localized regulation of inflammation (38). Here we reveal complex regulatory biology of Tnf, providing evidence that membrane-bound Tnf functions as a negative regulator of NF-κB likely through its binding to p75. Our results also support a model in which membrane-Tnf can function in an autocrine manner, acting within the cell that produces it. In this study, we bring together decades of research on Tnf biology and its role in inflammation using this single screening approach. Anti-Tnf therapy remains one of the most effective methods for the treatment of various autoimmune diseases, including rheumatoid arthritis (RA) and irritable bowel disease (IBD) (39,40), yet as many as 20-40% of patients don’t respond to treatment (41). Our findings show that editing of Tnf resulted in elevated Il6 levels, which is another important pro-inflammatory cytokine and might explain lack in response to anti-Tnf therapy. Kohanawa *et al* demonstrated that a Tnf -/- knock-out mouse showed elevated levels of Il6, which is consistent with our findings (42). More importantly, Kohanawa *et al* presented data showing production of TNF and IL6 can be negatively regulated by each other (42). Therefore, the targeting of Tnf could lead to elevated levels of Il6, which could result in elevated inflammation in patients. Our data showed that editing p55 inhibits NF-κB-driven Il6 production (Figure 4D-E). Blocking p55 (via antibody or small molecule) could be an alternative that could block the pro-inflammatory affects of p55, while allowing Tnf to instead bind to p75 to promote anti-inflammatory signaling (43). In summary, with one pooled CRISPR screen we have revealed interesting complexities of Tnf regulation and have presented evidence for alternative therapeutic strategies.

### Future directions

Here we have demonstrated the power of CRISPR screening in revealing important macrophage biology probing both viability and inflammatory pathways. Macrophages are critical cells of the innate immune system providing one of the first lines of defense against invading microbes. The ability for macrophages to function optimally requires their ability to proliferate and migrate in order to reach the site of infection and appropriately engage their inflammatory program. Here we found 60 macrophage-specific viability genes, as well as uncovering 115 novel regulators of NF-κB. The future characterization of these novel genes will likely yield novel regulatory insights into the complex regulation of viability and inflammatory pathways. From a therapeutic point of view, it would be interesting to explore whether there are drugs that may target some or any of our screen candidates (44). In conclusion, we have revealed important biology insights related to macrophage function and believe that this work represents a significant resource for the macrophage research community.

## Materials and Methods

### Cell lines

We have previously described the immortalized bone-marrow-derived macrophages with the NFKB reporter and Cas9 (iBMDM-NFKB-Cas9) cells (20). Cas9 activity was validated via GFP kd (~75% kd) prior to beginning the screen. Cells were cultured in DMEM, supplemented with 10% low-endotoxin fetal bovine serum (thermofisher) and 1X penicillin/streptomycin.

### sgRNA library design and cloning

We created genome-scale sgRNA library consisting of over 270,000 total sgRNAs (12 sgRNAs per gene) targeting every RefSeq-annotated (mm9) coding gene, as well as all microRNAs and select 3’UTRs. The library contains >5,000 non-target control sequences (NTC). The earliest possible “constitutive” exon of each transcript variant was targeted. The criteria for sgRNA selection and the cloning strategy protocol have been previously described (21,45). All sgRNA sequences are shown in Supplementary Table 1.

### Lentiviral production

HEK293T cells were seeded at 6,000,000 cells cells per plate in 15 cm dishes in 20 mL media (DMEM, 10% FBS) and incubated overnight at 37 °C, 5% CO2. The next morning, 8 µg sgRNA library plasmid, 4 µg psPAX2 (Addgene #12260), 4 µg pMD2.G (Addgene #12259) and 80 µL lipofectamine2000 (Invitrogen) were mixed into 1 mL serum-free OptiMEM (Gibco), vortexed and incubated for 20 min at RT and added to the cells. At 72 h post-transfection, supernatant was harvested, passed through 0.45 um filters (Millipore, Stericup) and aliquots were stored at −80 °C.

### CRISPR Screen

iBMDM-NFKB-Cas9 cells were infected with the sgRNA genome-scale library at a at low multiplicity of infection (moi=3) with an initial library coverage of >100X. Three days post infection, cells were puromycin-selected (10ug/ml) for 5 days obtain cherry-positive (sgRNA) cells and were maintained at >1000X coverage at all times.

### Growth screen

Prior to puro-selection, we collected a day 0 timepoint, consisting of 1000X coverage (270 millions cells). We then collected a day 20 timepoint (also 1000X coverage). Cells from both timepoints were cryo-preserved in 90% FBS, 10% DMSO for later processing.

### FACS screen

Library infected and selected iBMDM-NFKB-Cas9 cells were expanded to 2000X coverage. Cells were stimulated with 200ng/ml of LPS for 24h to induce expression of GFP (NFKB responsive). Prior to sorting, cells were collected in FACS buffer (1XPBS, 1%FBS, 5mM EDTA). Stimulated cells were FACS along side unstimulated cells to ensure the mean fluorescence intensity (MFI) of stimulated cells was >10-fold compared to unstimulated cells. All flow cytometry experiments and screening were conducted on a BD FACSAria II. GFP was excited using a 488-nm laser and detected using a 525/50-nm filter. Sorting was conducted using 4-way purity into 2 tubes and an 100-µm nozzle. Cells were gated by forward (FSC-A) and side scatter (SSC-A) for live cells, then for single cells using FSC-A/FSC-H. Lastly, we evaluated GFP expression (SSC vs GFP), FACS sorted and collected the top/bottom 20% into separate tubes. At least 100 cells/sgRNA (100X coverage) for each sorted population were collected and cryo-preserved in 90% FBS, 10% DMSO for later processing.

### gDNA processing, PCR and sequencing

Genomic DNA was collected from cell pellets (270 millions cells, 1000X coverage) or (27 millions cells, 100X coverage for sorted cells) and was extracted by methods described previously (21,45). A nested PCR strategy was used to 1) allow amplification sgRNA repertoire and 2) to add appropriate Illumina adapters for NGS (detailed protocol is described in (21)). For the 100X coverage sorted samples, we scaled the gDNA extraction volumes 1:5 (i.e. 2ml instead of 20ml). Quality and purity of the PCR product were assessed by bioanalyzer (Agilent), and sequencing was performed on an Illumina HiSeq 2500 platform using paired end 50 kits with the custom sequencing primer 5’-GAGACTATAAGTATCCCTTGGAGAACCACCTTGTTGG-3′ for reading the sgRNA sequence. Data was submitted to GEO.

### Screen Analysis and generation of hit list

fastq.gz files were analyzed using the gRNA_tool: https://github.com/quasiben/gRNA_Tool. All guide RNA (sgRNA) + barcode reads were collapsed to obtain raw sgRNA counts. Counts were normalized to the median and fold-changes were calculated for each sgRNA. To identify significant genes for both the growth screen and the FACS screen, the Mann-Whitney U test was performed comparing fold-changes for gRNAs targeting each gene to non-targeting controls (described in (46)). For the growth screen, the Day 20 sample was compared to the Day 0 sample. For the FACS screen, GFP low (bottom 20%) sorted cells were compared to GFP high (top 20%) sorted cells.

### gRNA selection for screen validation

For the 3’utr-targeting guides, we selected guides with >3-fold positive enrichment (Day20 vs Day0) that targeted the 3’utrs of essential genes with significant negative enrichment. The criteria for our NFKB candidate selection was as follows: 1) *Most significant:* We selected candidates with the lowest Mann-Whitney U test p-value. 2) *Novelty:* We focused on “novel” genes, which were not previously known to be involved in NFKB signaling. We used several databases including KEGG, String-DB and Cell Signaling TLR-signaling gene list to determine novelty of gene. 3) *Expression:* We evaluated expression and confirmed >10 FPKM for either un-stimulated or LPS-stimulation conditions. 4) *Viability:* We confirmed that our candidates did not have a significant viability phenotype. We targeted a total of 18 coding genes selecting guides with the strongest enrichment (2 guides/gene). We targeted microRNAs that showed >3-fold positive enrichment for at least 2 gRNAs. For positive controls we used guides targeting Tlr4

### 3’-UTR guide validation (Mix-cell growth assay)

gRNA-infected cells (cherry-pos) were mixed with uninfected cells (cherry-neg) at a 1:1 ratio in triplicate. We used FACS to monitored the ratio of cherry-pos to cherry-neg cells at 0, 10, 20 days post plating. All validation FACS was performed on the Attune NxT Flow Cytometer.

### NFKB guide Validation (qRT-PCR)

iBMDM-NFKB-Cas cells infected with indicated guide-expressing lentivirus and were stimulated with LPS (200ng/ml) for 6h prior to harvesting for RNA. Total cellular RNA from BMDM cell lines was isolated using the Direct-zol RNA MiniPrep Kit (Zymo Research) according to manufacturer’s instructions. RNA was quantified and controlled for purity with a nanodrop spectrometer. (Thermo Fisher). For RT-qPCR, 500-1000 ng were reversely transcribed (iScript Reverse Transcription Supermix, Biorad) followed by RT-PCR (iQ SYBRgreen Supermix, Biorad) using the cycling conditions as follows: 50C for 2 min, 95C for 2 min followed by 40 cycles of 95C for 15 s, 60C for 30 s and 72C for 45 s. qRT-PCR primer sequences are list below.

### ELISA Analysis

For the ELISA, iBMDMs were stimulated with LPS for 24h and supernatant was collected from triplicate wells. Supernatant was diluted 1:3 and Tnf-alpha levels were measured using the mouse TNF-alpha DuoSet ELISA (R&D Systems) kit following manufacturer’s protocol.

### Antibody staining for FACS

For intracellular staining, iBMDMs were LPS-stimulated for 0 or 6h and were treated with Brefeldin A for the last 5 hours of stimulation. Cells were then collected, fixed with 4% PFA and permeabilized with perm buffer (3% BSA, 0.2% Triton-X, 1XPBS), followed by antibody staining with PEcy7 anti-mouse Tnf-alpha (1:150, Thermofisher) or isotype control (1:150, Biolegend). For surface staining, iBMDMs were LPS-stimulated for 0, 6, 24h. Cells were collected in sorting media (2% Fetal Calf Serum, 5mM EDTA, 1XPBS), treated with Fc receptor block (1:250, BD Pharmingen) and were then stained with same Tnf and isotype antibodies used above, all done in sorting media.

### qRT-PCR primers

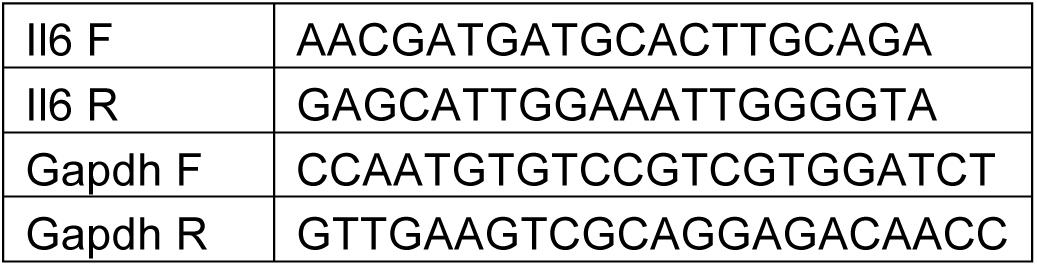

### RNA isolation and cDNA synthesis and RT-qPCR

Total cellular RNA from THP1 cell lines was isolated using the Direct-zol RNA MiniPrep Kit (Zymo Research) according to manufacturer’s instructions. RNA was quantified and controlled for purity with a nanodrop spectrometer. (Thermo Fisher). For RT-qPCR, 500-1000 ng were reversely transcribed (iScript Reverse Transcription Supermix, Biorad) followed by RT-PCR (iQ SYBRgreen Supermix, Biorad) using the cycling conditions as follows: 50°C for 2 min, 95°C for 2 min followed by 40 cycles of 95°C for 15 s, 60°C for 30 s and 72°C for 45 s. The melting curve was graphically analyzed to control for nonspecific amplification reactions.

### RNA-Sequencing

For generation of RNA-Sequencing libraries the human THP1 cells, RNA was isolated from control or Pam3CSK40-stimulated cells as described above and the RNA integrity was tested with a BioAnalyzer (Agilent Technologies). For RNA-Sequencing target RIN score of input RNA (500-1000ng) usually had a minimum RIN score of 8. RNA-Sequencing libraries were prepared with TruSeq stranded RNA sample preparation kits (Illumina), depletion of ribosomal RNA was performed by positive selection of polyA+ RNA. Sequencing was performed on Illumina HighSeq or NextSeq machines. RNA-seq 50bp reads were aligned to the human genome (assembly GRCh37/hg19) using TopHat. The Gencode V32 gtf was used as the input annotation. Differential gene expression specific analyses were conducted with the DESeq R package. Specifically, DESeq was used to normalize gene counts, calculate fold change in gene expression, estimate p values and adjusted p values for change in gene expression values, and to perform a variance stabilized transformation on read counts to make them amenable to plotting.

### Data sharing

All sequencing data generated from CRISPR screens and RNA-seq reported in this paper have been deposited to GEO under the ID code GSE138788.

## Acknowledgments

Thanks to all members of the Carpenter Lab and to Kate Fitzgerald and Maninjay Atianand for feedback on manuscript. This work was funded by an NIH R03 AI131019-01 and R21 AI142165.

Authors declare no conflict of interest.

---

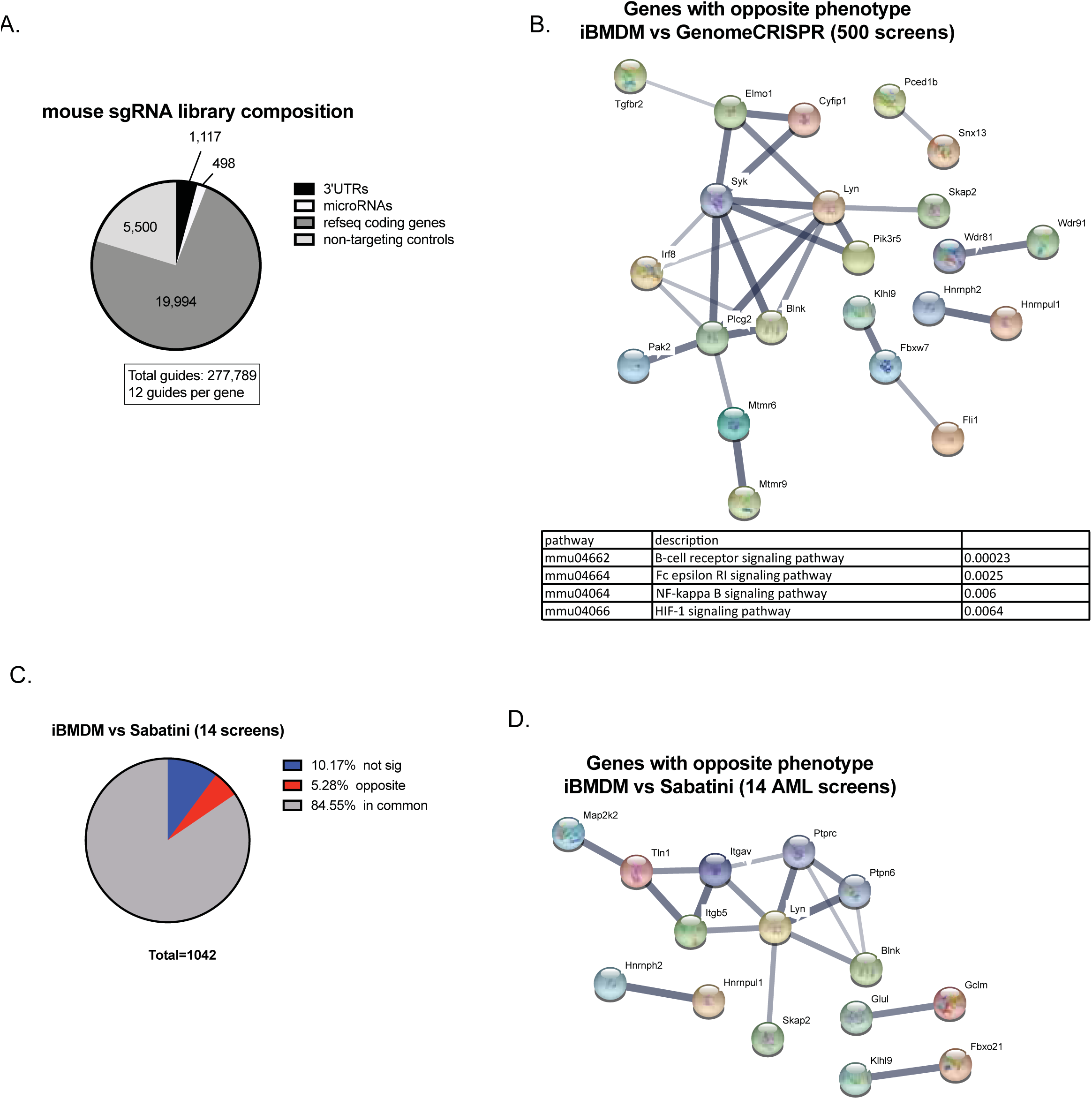

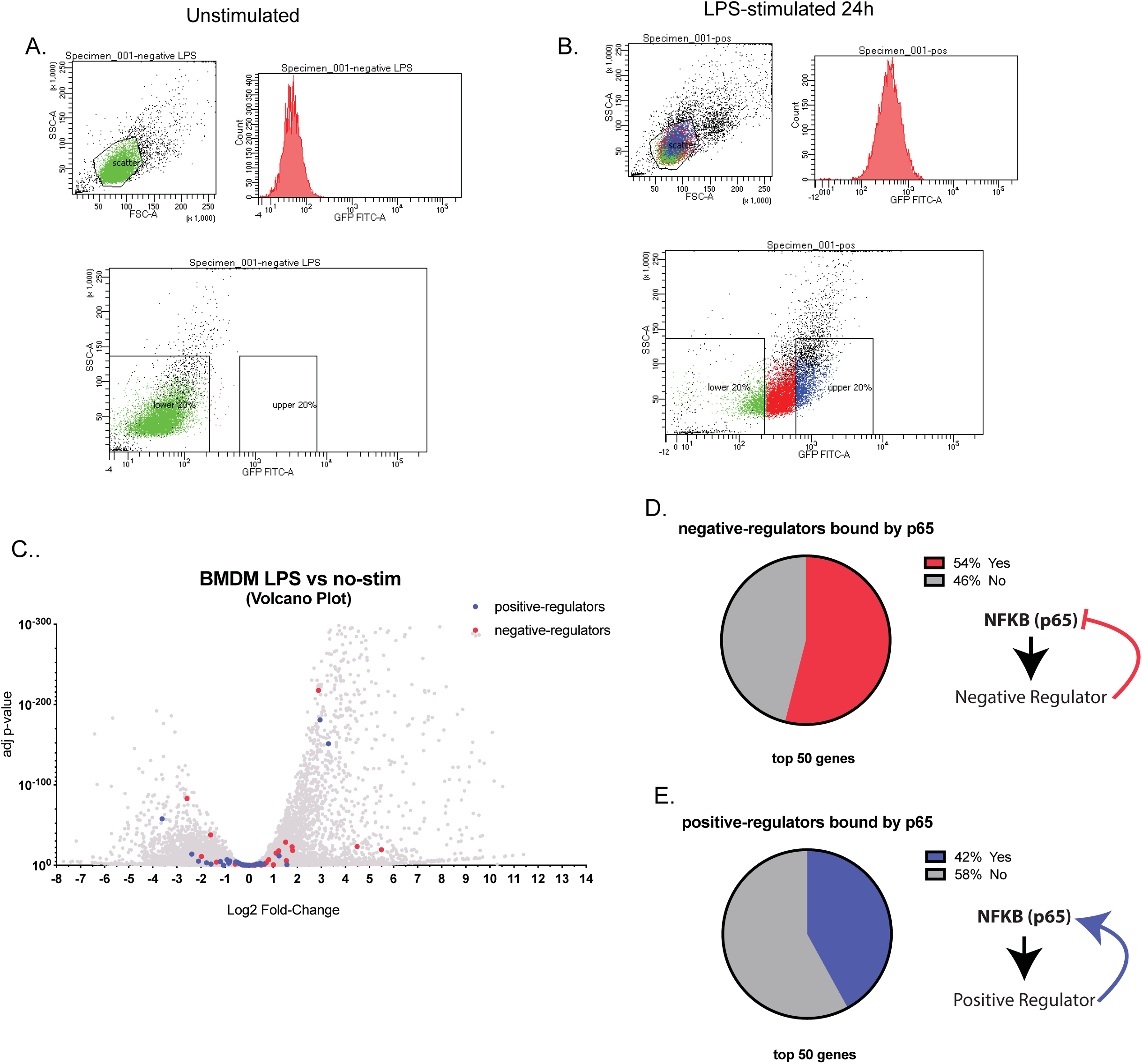

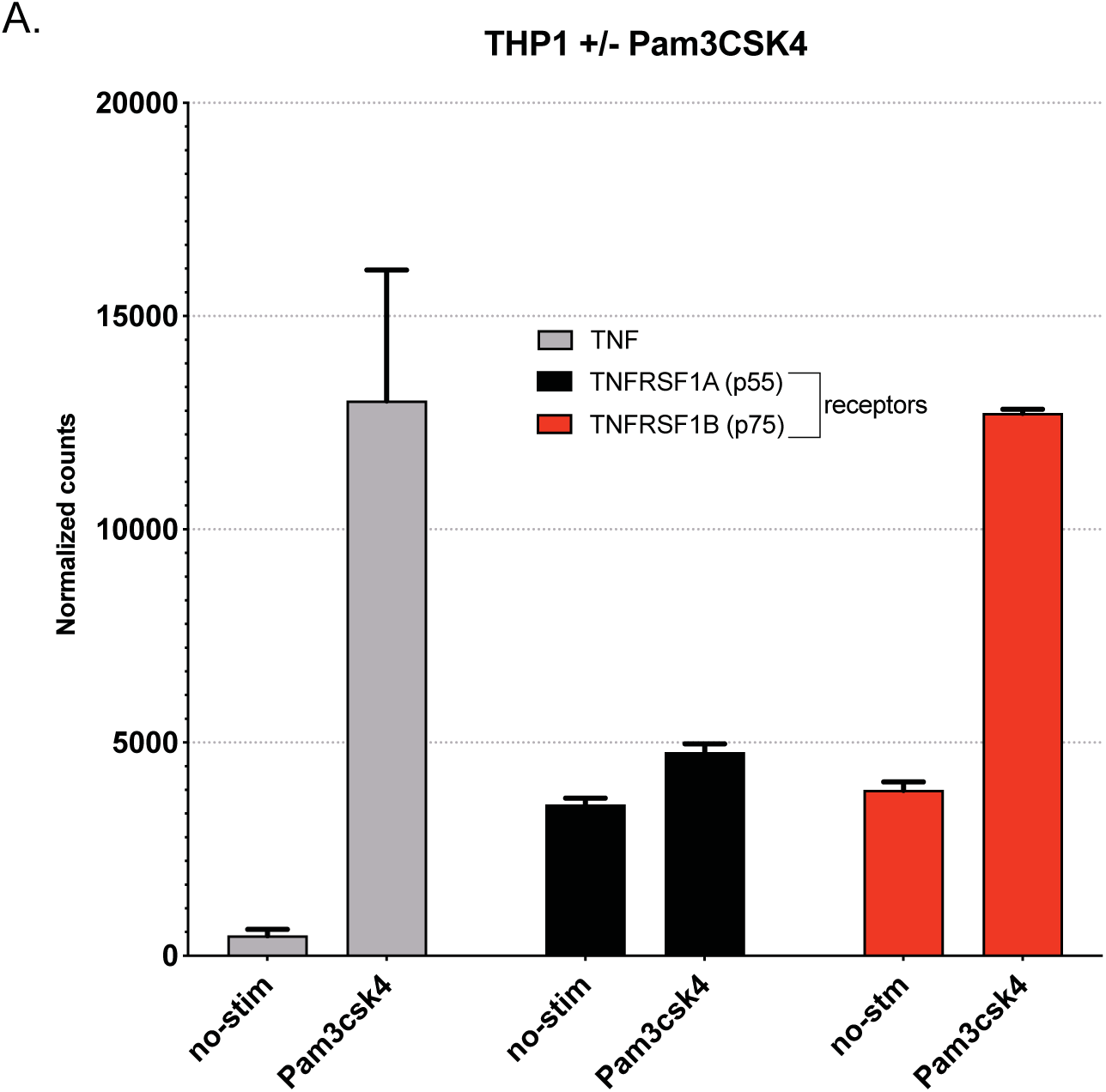

