## Supplemental Figure Legends for "High throughput CRISPR screening identifies genes involved in macrophage viability and inflammatory pathways"

**Figure S1: Comparison of current screen to CRISPR screen database.** A. Breakdown of all genes targeted by our custom mouse sgRNA library is displayed. B. Genes with opposite phenotypes in our screen compared to the GenomeCRISPR database are displayed visually using String-DB. KEGG pathway Go-term enrichment is also shown. C. Viability screen hits from our screen were compared to the Sabatini Lab ~14 Acute Myeloid Leukemia (AML) cell line CRISPR screens obtained from GenomeCRISPR (www.genomecrispr.org/). Genes "in common," genes showing "opposite" phenotypes and gene "not significant" are displayed. D. Genes with opposite phenotypes in our screen compared to the Sabatini Screens are displayed visually using String-DB.

**Figure S2: Expression and p65-binding of NFKB screen hits.** A-B. NF-κB FACS screen gating strategy for unstimulated cells (A) or 24h LPS stimulated cells (B). C. Differentially expressed genes in 6h LPS vs un-stimulated BMDMs is displayed as log2 fold-change vs. adjusted p-value volcano plot from previously published data (1). Expression of top 50 positive regulators (blue) and top 50 negative regulators (red) is shown . D-E. All p65 targets were determined using the ChIP-seq data from (2). Positive p65 binding was called if p65 peak was greater than 10 and was within 1kb of the annotated transcription start site (TSS). P65 promoter binding was then assessed for the top negative (D) and positive (E) regulators.

**Figure S3: Expression of TNF and TNF receptors in human cells.** A. Differentially expressed genes were determined for 6h Pam3CSK4-stimulated or un-stimulated THP1 (ATCC). Normalized counts +/- SD are displayed for TNF, TNFRSF1A and TNFRSF1B.

1. Zhang X, Chen X, Liu Q, Zhang S, Hu W. Translation repression via modulation of the cytoplasmic poly(A)-binding protein in the inflammatory response. Elife. eLife Sciences Publications Limited; 2017 Jun 21;6:619.

2. Lam MTY, Cho H, Lesch HP, Gosselin D, Heinz S, Tanaka-Oishi Y, et al. Rev-Erbs repress macrophage gene expression by inhibiting enhancer-directed transcription. Nature. Nature Publishing Group; 2013 Jun 27;498(7455):511–5.
